# Lipidomic profiling of clinical prostate cancer reveals targetable alterations in membrane lipid composition

**DOI:** 10.1101/2020.10.27.356634

**Authors:** Lisa M. Butler, Chui Yan Mah, Jelle Machiels, Andrew D. Vincent, Swati Irani, Shadrack Mutuku, Xander Spotbeen, Muralidhararao Bagadi, David Waltregny, Max Moldovan, Jonas Dehairs, Frank Vanderhoydonc, Katarzyna Bloch, Rajdeep Das, Jurgen Stahl, James Kench, Thomas Gevaert, Rita Derua, Etienne Waelkens, Zeyad D. Nassar, Luke A. Selth, Paul J. Trim, Marten F. Snel, David J. Lynn, Wayne D. Tilley, Lisa G. Horvath, Margaret M. Centenera, Johannes V. Swinnen

## Abstract

Dysregulated lipid metabolism is a prominent feature of prostate cancer that is driven by androgen receptor (AR) signaling. Herein, we used quantitative mass spectrometry to define the “lipidome” in prostate tumors with matched benign tissues (n=21), independent tissues (n=47), and primary prostate explants cultured with a clinical AR antagonist, enzalutamide (n=43). Significant differences in lipid composition were detected and spatially visualized in tumors compared to matched benign samples. Notably, tumors featured higher proportions of monounsaturated lipids overall and elongated fatty acid chains in phosphatidylinositol and phosphatidylserine lipids. Significant associations between lipid profile and malignancy were validated in unmatched samples, and PL composition was characteristically altered in patient tissues that responded to AR inhibition. Importantly, targeting of altered tumor-related lipid features, via inhibition of acetyl CoA carboxylase 1, significantly reduced cellular proliferation in tissue explants (n=13). This first characterization of the prostate cancer lipidome in clinical tissues revealed enhanced fatty acid synthesis, elongation and desaturation as tumor-defining features, with potential for therapeutic targeting.

## Introduction

With more than 1 million deaths annually, prostate cancer remains a major cause of mortality and morbidity for men worldwide (1). The clinical implementation of systemic androgen receptor (AR) targeting agents such as enzalutamide and apalutamide has increased the available therapeutic options beyond androgen deprivation, but development of resistance to these strategies remains inevitable. To more effectively combat this disease, there is a need for alternative targets for intervention and a thorough understanding of the molecular changes that accompany cancer development, progression and therapy. The advent of parallel ‘omic’ approaches has led to the identification of previously unsuspected cancer subtypes and therapeutic targets. However, in contrast to the genome, transcriptome, and proteome, the cancer “lipidome” remains inadequately characterized (2). As building blocks of cellular membranes, lipids affect numerous cellular processes including signal transduction, ion transport, cell proliferation, energy metabolism and cell death mechanisms, which are all involved in the development and progression of cancer (3,4).

For prostate cancer, the lipidomic changes that accompany malignancy are of particular interest due to the unique metabolic profile of this cancer, whereby the normal cellular production of citrate is instead utilized in the TCA cycle for oxidative phosphorylation and biosynthetic processes such as lipogenesis (5,6). Moreover, lipid metabolism is a highly androgen-sensitive process in prostate cancer cells (7) and lipid composition may therefore be a unique cellular readout of both androgen targeting and tumorigenesis. While panels of circulating plasma lipids have previously been associated with prostate cancer risk (8), diagnosis (9) and patient outcome (10), analysis of the prostate tumor lipidome has largely been confined to cell line-based studies (11–13), which lack clinical relevance. The recent evidence of malignancy-related changes in lipid composition of prostate tumors provided by mass spectrometry-based imaging studies (14–17) supports undertaking a more detailed and quantitative study of the clinical prostate cancer lipidome. Moreover, treatment-related changes in the lipidome, which may reveal new resistance-related vulnerabilities, remain completely unexplored.

To gain insight into the potentially targetable changes in the lipid composition of prostate cancer, we employed a quantitative mass spectrometry-based lipidomics approach coupled with mass spectrometry imaging to robustly analyze and visualize a wide range of intact phospholipid (PL) species in malignant and matched non-malignant tissues. Our results provide the first comprehensive picture of the lipidomic landscape in a clinical cancer context and reveal robust associations with malignancy. We further demonstrate treatment-related changes in the lipidome accompanying response to enzalutamide, an AR antagonist widely used in the management of prostate cancer, in patient-derived explants (PDEs) of clinical prostate tissues. Finally, we report that an inhibitor of acetyl CoA carboxylase was effective in pharmacologically targeting the most recurrent lipidomic alterations observed.

## Results

### Tumor-specific lipid profiles are evident in clinical prostate tumors

Spatial variation in PL composition was initially assessed in a set of clinical prostate tumors that contained discrete benign and malignant areas of epithelium within the same tissue section, using MALDI-mass spectrometry imaging (MALDI-MSI). Shown in ***Figure 1A*** is a representative tumor in which comparison of spectral data from these regions revealed a tumor-specific PL composition that was distinct from that detected in pathologically benign epithelium or a region of high-grade prostatic intraepithelial neoplasia (PIN) from the same patient (***Figure 1B***; Supplementary Figure 1A). Distinct PL mass spectra and specific masses (eg m/z 756.53, ***Figure 1A***) were consistently detected in multiple independent tumor regions assessed across 3 individual PCa patients (Supplementary Figure 1B,C), consistent with recent reports (16,17) that characteristic changes in lipid composition accompany prostate tumorigenesis.

**Figure 1.**
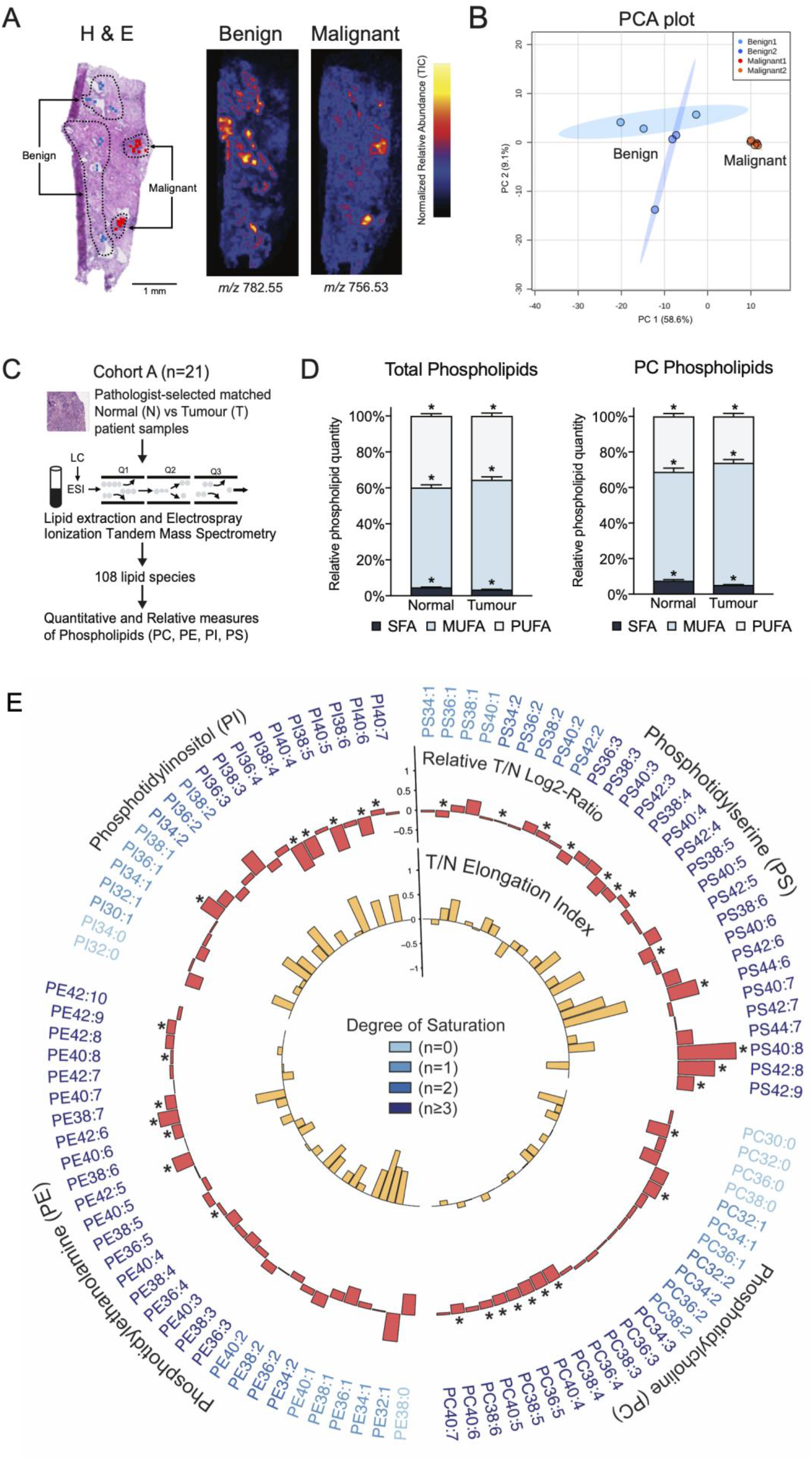
Evidence for a tumor-associated phospholipidome in clinical prostate cancer. **A.** MALDI-mass spectrometry imaging of pathologically heterogeneous prostate tissue, including ion maps of two representative examples of histology-restricted lipid masses. **B.** principal component analysis of the top 25 mass features distinguishing benign from malignant regions of tissue. **C.** Workflow for shotgun lipidomics analysis of matched normal and tumor tissues from 21 prostate cancer patients. **D.** Relative proportions of phospholipids containing 0 (SFA), 1 or 2 (MUFA), or 3 or greater (PUFA) unsaturations in benign versus tumor specimens. **E.** Circle plot of tumor-related lipidomic changes in relative abundance and fatty acyl chain elongation across the patient cohort. The outer circle represents the ratio of median-adjusted normalized abundance of individual lipid species in tumor versus benign tissues. * reflects significant association with tumor tissues. The inner circle represents fatty acid elongation index in tumor versus benign tissues. PL are annotated using “lipid subclass” followed by the “total fatty acyl chain length:total number of unsaturated bonds”.

In light of these findings, we undertook a more detailed examination and quantification of tumor-related changes in PL composition using shotgun lipidomics incorporating electrospray tandem mass spectrometry in pathologist-microdissected regions of prostate tumor and matched benign tissues from 21 prostate cancer patients (***Figure 1C***). Clinicopathological data for the patients are summarized in Supplementary Table S1. Using this methodology, a total of 108 PL species of the four most abundant subclasses (PC, PE, PS and PI) could be detected and quantified. PL profiles of both cancer and normal tissues were dominated by PC species, and the total proportion of PC lipids was greater in tumor than in benign tissues for this cohort (Supplementary Figure 2A; p<0.05). There was a significant tumor-specific increase in the collective abundance of PL species with 1 or 2 double bonds, most evident in the PC lipids (***Figure 1D***). As these mainly represent species with one or two monounsaturated fatty acyl (MUFA) chains, this shift altered PL composition towards a greater overall proportion of MUFAs in the tumors, consistent with a lipogenic tumor phenotype (11).

Shown in ***Figure 1E*** is a circle plot summarizing the individual tumor-related lipidomic changes for this cohort of patients. PL are annotated by “lipid subclass” followed by the “total fatty acyl chain length:total number of unsaturated bonds” (eg PC34:1). The species in each PL class are ordered from fully saturated to highly polyunsaturated and, within each subgroup, from shortest to longest (combined) acyl chain length. When considering the relative abundance of individual PL species, consistent patterns of change in PL saturation groups were more evident within certain lipid classes (***Figure 1E***, outer circle). Most notably, PC species had tumor-related increases in fatty acyl chains containing 1 or 2 double bonds, indicative of MUFAs, accompanied by relative decreases in polyunsaturated (PUFA; ≥3 double bonds) and fully saturated (0 double bonds) species. In PS species, however, marked increases in long PUFA species were evident across multiple saturation groups in tumor compared to benign tissue.

Consistent changes across patients were also detected in the fatty acyl chain lengths of tumor PLs. To most optimally visualize altered acyl chain length, we expressed the abundance of each PL species in tumors and matching benign tissue relative to the shortest PL species of each saturation subclass (***Figure 1E***, inner circle; denoted “elongation index”). Whereas substantial heterogeneity in fatty acyl chain length existed between individual patients for PC and PE lipids (Supplementary Figure 2B), almost every tumor exhibited increased combined acyl chain length over multiple saturation groups of the PI and PS classes compared to normal tissue, particularly for polyunsaturated species (Figure 1D; Supplementary Figure 2B). This was validated within the cohort as a significant increase in average chain length for polyunsaturated PI and PS lipids (Supplementary Figure 2C).

### Associations between Lipid Profiles and Malignancy

Based on the above observations, we investigated whether there were associations between any lipid measures and sample malignancy status. Linear mixed effects models regressed PL measures from malignant samples onto the equivalent values from benign samples, adjusting for age and batch. We specifically considered i) abundance of individual PL species, ii) saturation group abundance and iii) fatty acyl chain length per saturation group. The permutation analysis indicated that there were associations between lipid profile and malignancy beyond what would be expected by chance (Supplementary Figure 3A; permutation p<0.001). Of the 51 lipid features most strongly associated with malignant versus benign prostate tissue, 34 were individual lipid species (***Figure 2A*** & Supplementary Table S2), five were mean chain lengths within saturation groups and 12 were overall saturation group abundance (Supplementary Figure 3B & Table S2). As expected, certain individual species crosscorrelated within and across head group classes (***Figure 2B***).

**Figure 2.**
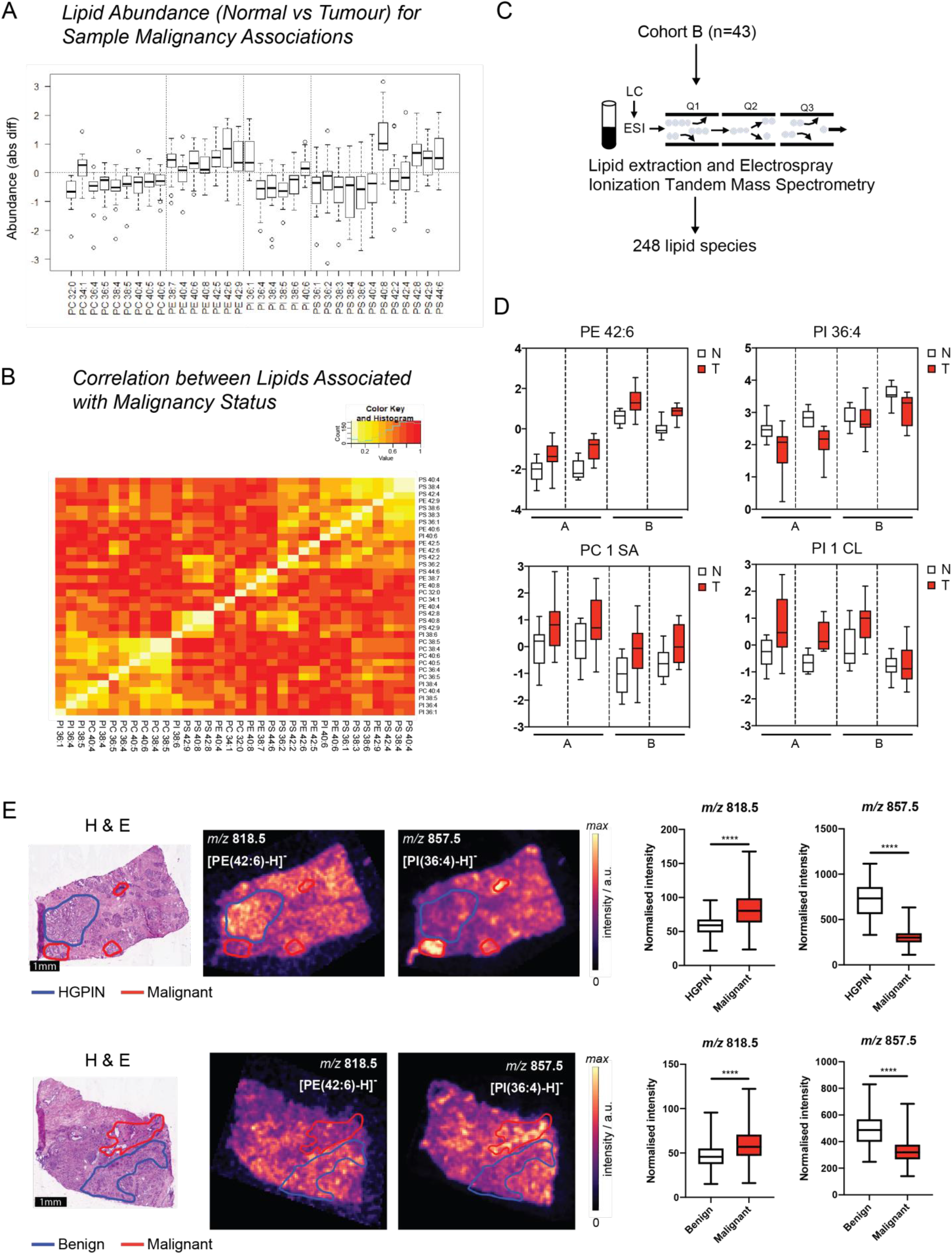
Associations of phospholipid profile with malignancy. **A.** Phospholipid variables significantly associated with tumor versus matched benign tissues in Cohort A. **B.** Correlation plot of individual phospholipid species significantly associated with sample malignancy in Cohort A. **C.** Workflow for lipidomics analysis of unmatched patient tissue cohort B (n=47). **D.** Box plots of representative examples of different phospholipid variables significantly associated with sample malignancy across both patient cohorts A and B. **E.** MALDI-mass spectrometry imaging of ion masses corresponding to PE42:6 (*m/z* 818.5) and PI36:4 (*m/z* 857.5) in 2 independent prostate cancer tissues, with box plots of normalized ion intensity between non-malignant and malignant tissues regions adjacent to the images.

We subsequently assessed whether similar malignancy-associated lipidomic profiles would also be evident in an independent collection of 47 non-patient matched tissue specimens comprising 26 tumors and 21 benign samples (***Figure 2C***). A greater range of lipids were measured in this cohort (n=248), and included sphingomyelins (SM), ceramides (Cer) and lysoPLs (containing only 1 fatty acyl chain) in addition to the main four PL classes analyzed above. Using the same criteria as for the paired cohort, a substantial signal was again detected in terms of FDR for significant associations between lipid features and sample malignancy (Supplementary Figure 3A). Associations were detected for 64 lipid features, including 44 individual lipid species (FDR=0.03; permutation p<0.001; Supplementary Figures 3C,D,E and Supplementary Table S3).

There was considerable overlap in the lipid features identified in both cohorts, and by combining data from both cohorts, we identified a series of lipids that robustly associated with prostate tumor malignancy (Table 1). Shown in ***Figure 2D*** are representative examples of these lipids demonstrating concordant associations with malignancy across the individual cohorts and batches, which included the abundance of monounsaturated PC lipids (PC1 SA), individual lipids PE42:6 and PI36:4, and chain length in monounsaturated PI lipids (PI1 CL). Consistent with these data, MALDI-MSI on two patient tissues imaged in negative ion mode revealed the expected changes in relative abundance of masses corresponding to PE42:6 (*m/z* 818.5) and PI36:4 (*m/z* 857.5) in malignant versus non-malignant regions of the tissues (***Figure 2E***; box plots of normalized mass intensity shown adjacent to ion map images; Supplementary Figure 4).

**Table 1.**
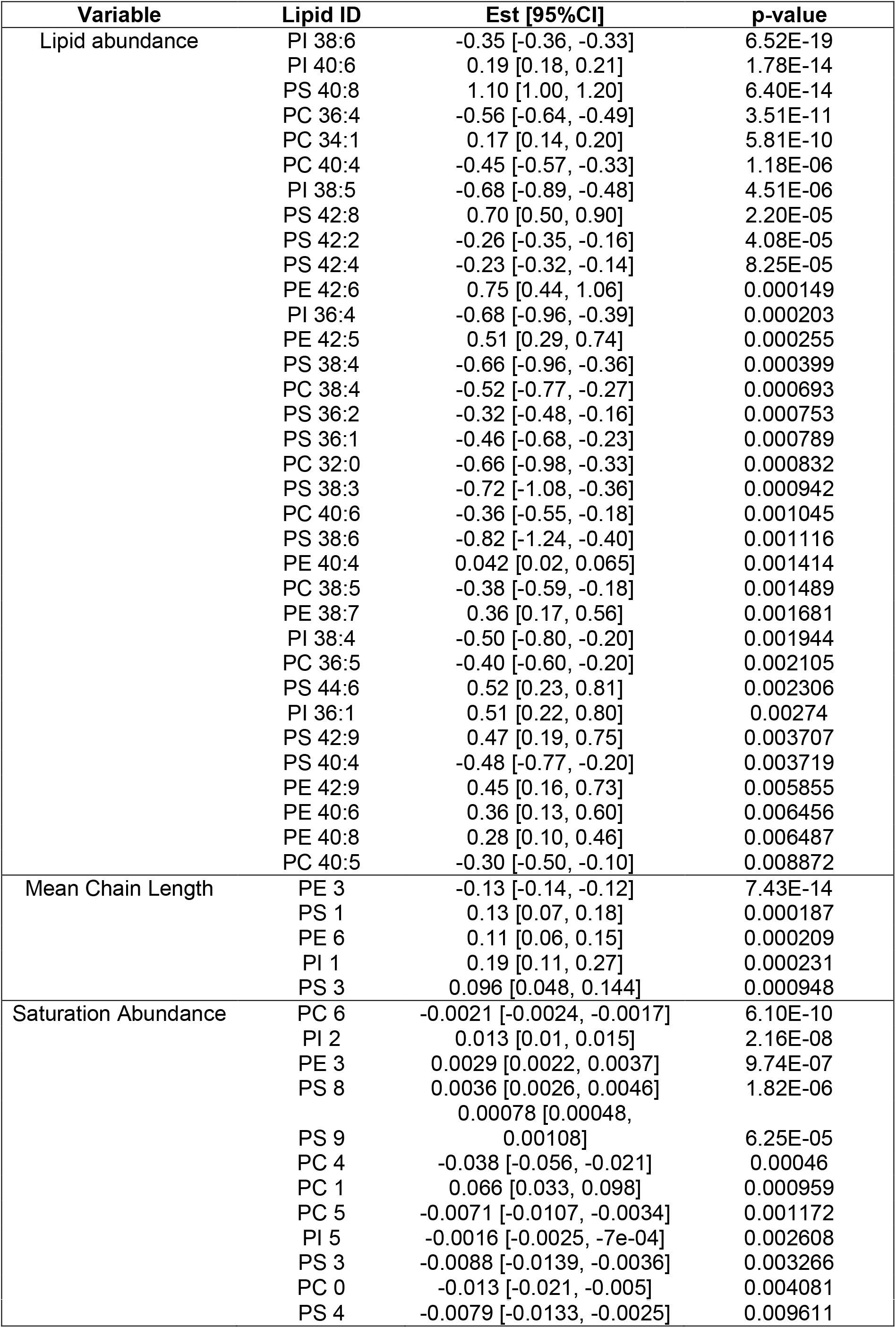
Conserved lipid associations with malignancy in clinical prostate samples.

While the tumor cohort size limited our ability to detect associations with clinical parameters such as serum PSA (Supplementary Figure 5B; permutation p=0.79) and Gleason Score (Supplementary Figure 5B,C; permutation p=0.24), there was evidence for weak associations between lipid profile and TMPRSS2-ERG subtype (Supplementary Figure 5B,D,E and Table S4; permutation p=0.05) and proliferation (Ki67 positivity index) in malignant samples (Supplementary Figure 5A,F and Table S5; permutation p=0.10).

### Lipid profile is altered by AR inhibition in patient-derived tumor explants

From the cohort of 47 unmatched samples analyzed above, 43 of the samples were also cultured *ex vivo* as patient-derived explants (PDEs) in the absence and presence of the clinical antiandrogen, enzalutamide (ENZ; n=24 cultured with 10μM ENZ, n=19 cultured with 10μM and 50μM ENZ; ***Figure 3A***). This provided the unique opportunity to examine dynamic, treatment-related changes in lipid composition using patient-matched samples. *Ex vivo* culture alone had a minimal effect on lipid profile; 80% of lipid species had a correlation >0.5 between uncultured and cultured tissues (Supplementary Figure 6A). Transcript profiling and gene set enrichment analysis performed on a subset of these samples (n=12) confirmed significant downregulation by ENZ (10μM) of androgen signaling and multiple pathways associated with metabolism (***Figure 3B***). Independent qPCR validation confirmed the decrease in expression of canonical AR target genes *kallikrein 3* (encoding prostate specific antigen) and *kallikrein 2* gene across the majority of samples (Supplementary Figure 6B). As expected based on clinical trial outcomes (18,19), the PDEs showed substantial heterogeneity in proliferative response to ENZ, measured as change in Ki67 proliferative index from matched vehicle-treated tissue (***Figure 3C***). The above features of the ENZ-treated PDEs were conducive to analysis of treatment-related dynamic changes in PL profile, and associations with decreased Ki67 index in individual samples. ENZ-related changes in 18 PL variables (including 15 individual species, featuring long-chain PC and PE lipids) were associated with Ki67 proliferation index (Supplementary Figure 6C and Supplementary Table S6; ***Figure 3D***). MALDI-MSI analysis of a subset of PDE tissues indicated that the majority of individual response-related lipids identified reflected the epithelial regions of the tumors, and treatment-related changes in abundance were evident (example of PC34:1 shown in ***Figure 3E***, boxplots in Supplementary Figure 6D). Dynamic changes in PL profile were therefore associated with response to AR inhibition across this cohort of PDEs.

**Figure 3.**
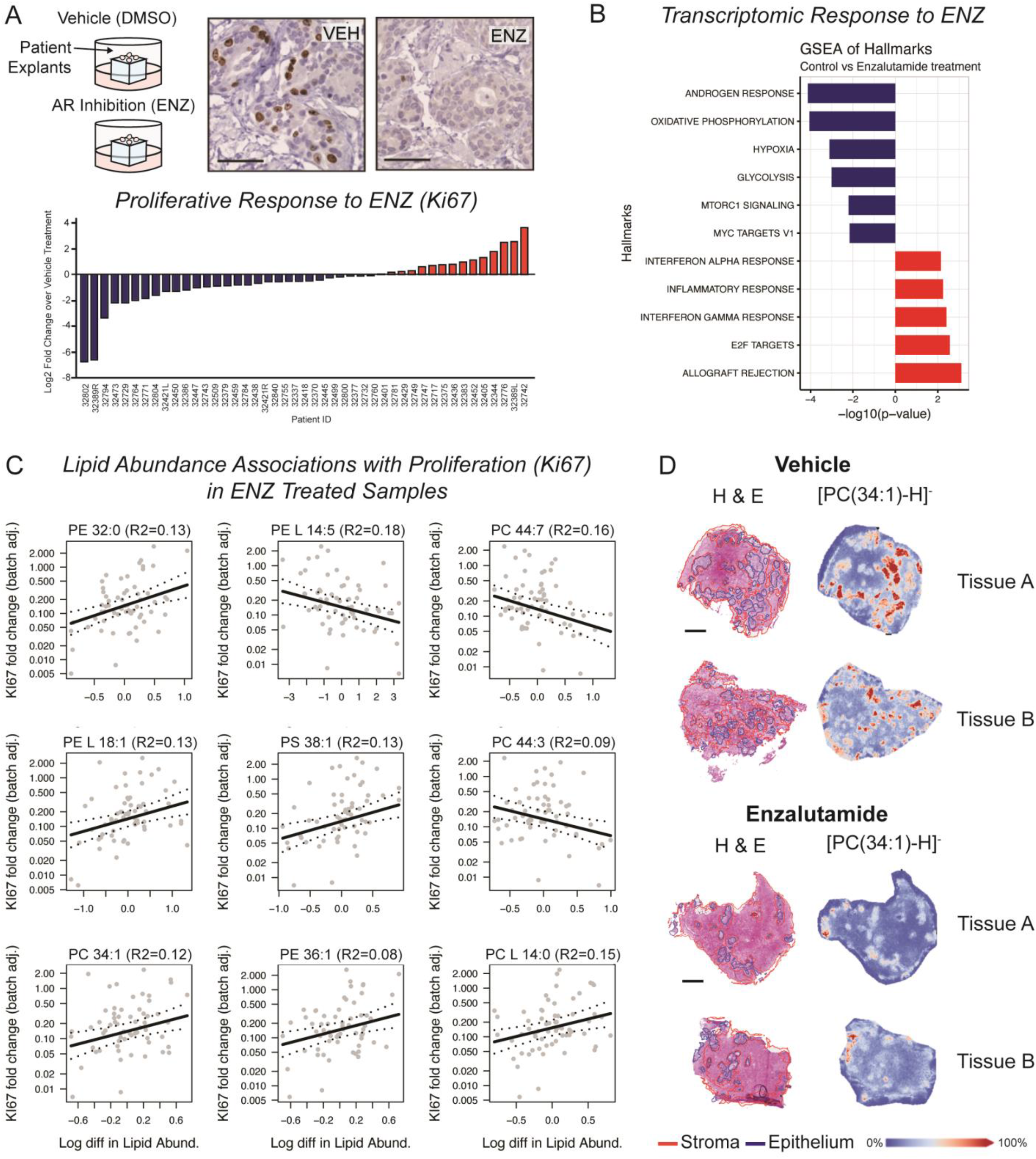
Associations between phospholipid profile and tumor response to the clinical antiandrogen enzalutamide (ENZ). **A.** Variation in proliferative response of prostate tissues to ENZ across cohort B. Upper panel: Patient-derived explant culture setup for prostate tissue from Cohort B and Ki67 immunohistochemistry in a representative set of patient samples. Lower panel: Waterfall plot of individual patient response to ENZ, measured as log2fold change in Ki67 proliferative index. **B.** Gene set enrichment analysis for transcriptomic data from enzalutamide treated versus vehicle control tissues (n=12 patients). **C.** Correlation plots of phospholipid species whose change in abundance is significantly associated with response (change in Ki67 index) to ENZ (p<0.01). **D.** MALDI-mass spectrometry imaging of ion mass corresponding to PC34:1-H+ in 2 independent prostate cancer tissue cores from a single patient, showing epithelial localization and ENZ-related change in abundance.

### Targeting lipidomic changes in clinical tumors suppresses cellular proliferation

To determine whether the cancer-associated alterations in PL composition directly influence cellular proliferation, or merely accompany tumorigenesis, we pharmacologically targeted two of the key lipidomic changes we detected (i.e. enhanced lipogenesis and elongation of fatty acyl chains in the PLs) in patient-derived explants (PDEs) of clinical prostate tissues. Inhibition of acetyl CoA carboxylase (ACC1/2), which depletes the cellular content of malonyl CoA, was selected as a strategy to simultaneously inhibit both synthesis and elongation of fatty acids (***Figure 4A***). We utilized PF-05175157, an ACC1/2 inhibitor (20), as a proof of principle tool. Culture of PDEs with PF-05175157 (50μM) for 48 or 72 hours (***Figure 4B***) markedly suppressed epithelial cell proliferation in the tumors (n=13) compared to matched, vehicle-treated control tissues (***Figure 4C***). Enhanced epithelial staining of pACC1, a marker of ACC inhibition by PF-05175157 in prostate cancer cells (Supplementary Figure 7B), confirmed that the agent was effectively targeting ACC activity in the tumors (p<0.01; ***Figure 4D***). Moreover, PL profiling by mass spectrometry revealed a pronounced shortening of PL fatty acyl chains in the majority of PF-05175157-treated tumors (***Figure 4E***; Supplementary Figure 7). Together, these data link the efficacy of PF-05175157 to ACC1 inhibition in the tissues.

**Figure 4.**
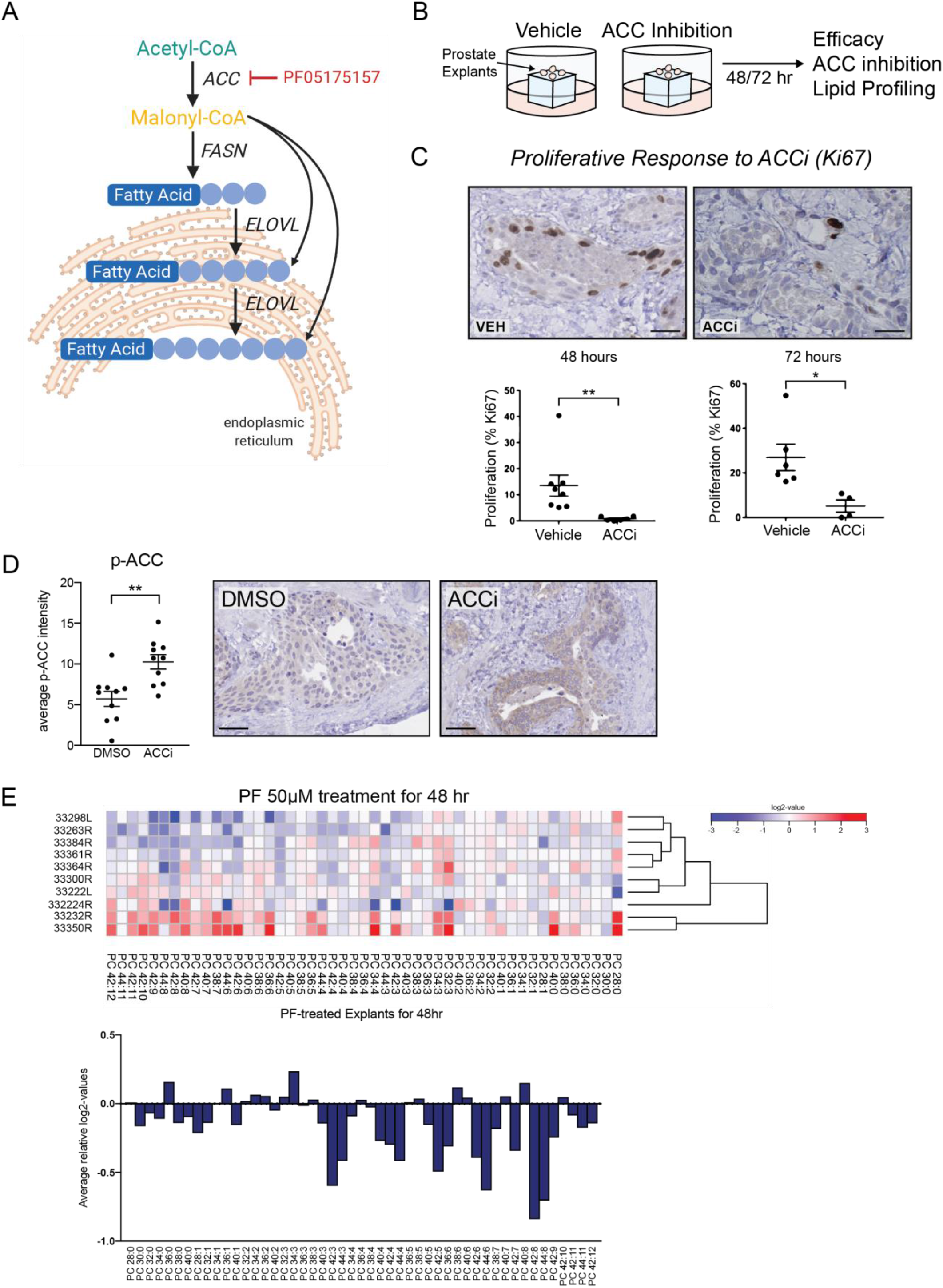
Efficacy of acetyl CoA carboxylase inhibition in patient-derived prostate explants. **A.** The ACC1/2 inhibitor PF-05175157 targets fatty acid synthesis and chain elongation. **B.** Patient-derived explant culture setup and workflow. **C.** Antiproliferative activity, measured by Ki67 proliferative index, of PF-05175157 (50μM) compared to vehicle-treated matched control tissue in patient-derived explants (PDEs; n=13 patients). **D.** Immunohistochemical detection and quantification of pACC1 intensity in PDEs cultured in the absence and presence of PF-05175157. **E.** Treatment-induced alterations (expressed as log2-fold change in the heatmap) in PC phospholipid abundance for PDEs cultured with PF-05175157. Altered relative fatty acyl chain length for the upper cluster of PDEs are further visualized graphically.

## Discussion

Using mass spectrometry-based lipidomics to sensitively quantify and visualize PL species in clinical tissues, we have provided new insight into the changes in lipid composition that accompany prostate cancer development. Moreover, this is the first report of dynamic, treatment-related changes in lipidomic profiles and the efficacy of targeting lipid metabolic enzymes in a clinical tissue context. This study therefore moves beyond previous cell line-based approaches to demonstrate that, despite the observed heterogeneity at the level of individual lipid species, recurrent and clinically-actionable changes in PL metabolism can be detected in tumors; some of which may represent common vulnerabilities. Lipidomic profiling of prostate tumor biopsies, possibly guided by imaging approaches as highlighted in several recent reports (16,17), has the potential to provide new information about disease features and, potentially, patient responsiveness to therapeutics such as enzalutamide.

While robustly associated with sample malignancy, PL profile was only weakly linked to the most common clinicopathological characteristics of prostate cancer; PSA levels and Gleason score. While this may reflect limitations in the sample size of our cohorts, it also raises the possibility that lipidomic profiling may provide independent information regarding tumor biology and prognosis. Moreover, the associations we detected between lipid profiles and the TMPRSS2-ERG molecular subtype and Ki67 proliferative index are interesting observations that warrant further investigation in larger independent tissue cohorts, particularly in light of our recent report of Ki67 status in localized prostate cancer being a significant predictive biomarker of subsequent metastatic relapse (21).

Despite differences in the breadth and scope of lipid classes measured, certain key cancer-related changes in PL profile reported here support the findings of previous imaging-based studies, notably for altered abundance and/or elongation of PC and PI-based lipids and ceramides (reviewed in (2,22)). While the functional consequences of individual lipid changes remain to be elucidated, PI lipids are of fundamental importance in cancer cells as they form the membrane scaffold for kinase and phosphatase activity that supports oncogenic signaling. Moreover, altered acyl chain length of PI-based lipids has been linked to p53 mutational status (23), a common genomic alteration in clinical prostate cancer. The results are also largely consistent with our earlier observed overexpression of lipid synthetic enzymes, such as FASN, in cancer cells, which correlates with a shift towards MUFA-containing species, at the expense of PUFA-containing species (11). Similarly, the tumor-related shift from saturated to monounsaturated PC species that we found to be significantly associated with sample malignancy is concordant with the earlier observed overexpression of SCD in certain cancer tissues (24–27). The novel changes in acyl chain length and head group switches that we detected may be related to reported alterations in enzymes involved in acyl chain elongation including ELOVL enzymes, modulators of malonyl-CoA levels including ACC, FASN and malonyl-CoA decarboxylase and head group-modifying enzymes including PS decarboxylase (28–32). Tumor-specific activation/inactivation patterns of these individual enzymes, most likely driven by tumor-specific oncogenic signaling (33–38), may lead to the unique phospholipid profile that is characteristic for every individual tumor. In view of the evidence that the lipid composition of cellular membranes affects numerous aspects of cell biology (reviewed in (39)), including membrane fluidity and curvature, vesicle formation, signal transduction (40), ion channel activity (41,42), susceptibility to lipid peroxidation (11), resistance to oxidative stress (11), energy metabolism (43), and uptake and response to chemotherapeutics (11), even subtle changes within the lipidome may be critical to support the cancer phenotype and treatment resistance. Given the lipidome is an integration of oncogenic events and an effector of numerous cancer-related processes, it is expected that the tumor lipidome holds significant potential for biomarker discovery and identification of novel targets that may be used in a theranostic setting (44).

While inter-patient heterogeneity was evident in PL profiles from clinical prostate tissues in the current study, particularly at the individual lipid species level, more consistent changes in broader lipid metabolic processes were evident. Notable among these were proportionally higher fatty acid monounsaturation, and elongation of the PL fatty acyl chains. Considering these phenotypes, inhibition of acetyl CoA carboxylase (ACC) presented an appealing approach to simultaneously target *de novo* biosynthesis and elongation of intracellular fatty acids by restricting production of the substrate for long chain fatty acid biosynthesis, malonyl CoA (45,46). To date, the focus for development of ACC inhibitors has been their ability to inhibit *de novo* lipogenesis and increase fatty acid oxidation, thereby reducing lipid accumulation and improving insulin sensitivity in patients with diabetes or non-alcoholic liver steatosis. There has, however, been considerable interest in repurposing these agents for oncology, particularly for lipogenic tumors such as prostate and breast, in which potential side-effects would be less of a concern. Here, as proof of principle, we utilized a spiroketone derivative ACC1/2 inhibitor, PF-05175157, developed by Pfizer as a clinical agent for treatment of Type 2 diabetes and non-alcoholic hepatic steatosis (20). Our results show marked efficacy for this agent in reversing the chain elongation phenotype and reducing epithelial cell proliferation in clinical PDEs, raising the promise that this class of agents may be efficacious in clinical prostate cancer. While the possibility of off-target effects of this compound contributing to its antiproliferative effects cannot be discounted, we have used two lines of evidence to associate efficacy with ACC inhibition in the tissues. First, we assessed the tissue levels of Ser79-phosphorylated ACC1, which is the inactive form of ACC1, the predominant isoform in prostate cells. This phosphorylation was induced by PF-05175157 treatment of prostate cancer cells and in PDEs. Second, our PL profiling of the treated tissues revealed consistent shortening of the fatty acyl chains, indicative of decreased elongation reactions. Taken together, our findings provide the first evidence that ACC inhibition can be achieved in the context of a complex tumor microenvironment and provide impetus for further investigation of ACC inhibition in prostate cancer. In light of the marked tumor-associated changes in fatty acyl chain saturation, and particularly monounsaturation, future studies targeting desaturases such as SCD1, shown recently to be a promising therapeutic target in prostate cancer (47), would also distinguish causality from association for this alteration.

In summary, the cancer-related changes in PL profiles we have detected in clinical tissues strengthen the case for lipidomics as a source of novel molecular cancer biomarkers and therapeutic targets, as well as an indicator of the underlying biology from which PL profiles are derived. Defining subtypes of lipid profile in tumors, rather than immortalized cell lines, and the underlying mechanisms in this and other association-based studies are all critical future endeavors if key components of lipid metabolism are to be effectively targeted in clinical disease. The heterogeneity of PCa evident from our work reinforces the concept that such approaches must also be personalized to the individual patient’s biology. Our findings warrant further functional investigation of lipidomes in other cancer types, to unravel the molecular mechanisms underlying these changes and to explore the impact on membrane functioning and, ultimately, on cancer development and progression.

## Materials and Methods

### Tissue collection

#### A. Matched normal/tumor cohort

Prostate tumor tissues with matching normal samples were obtained from patients who had undergone radical prostatectomy (Centre Hospitalier Universitaire de Liège, Belgium). Samples were snap-frozen and stored at −80°C for lipid and protein extractions. Normal and tumor tissues were identified by histological analysis of adjacent tissue, Gleason scores were determined and the percentage of cancer was estimated (48). All tumor samples used for lipidomics were verified to contain at least 75% prostate adenocarcinoma by histological examination. The Local Commission for Medical Ethics and Clinical Studies at the University of Liège approved the use of clinical samples. Approval to perform lipidomics analysis on clinical samples was obtained from the local Ethical Committee of KU Leuven.

#### B. Unmatched patient tissue cohort

Prostate tissues were collected with written informed consent from patients undergoing radical prostatectomy at St Andrew’s Hospital, Adelaide, Australia. A longitudinal section of each tissue was removed prior to *ex vivo* culture (described below). Half was snap frozen and the remainder fixed in formalin and paraffin embedded for assessment by a pathologist. Ethical approval for tissue collection and experimentation was obtained from St Andrew’s and the University of Adelaide Human Research Ethics committees. Histopathologic features of all tumors used in this study are detailed in Supplementary Table 1; an additional 5 patients with only benign prostatic hyperplasia and no evidence of cancer were also analyzed with this cohort.

### *Ex vivo* culture of primary prostate tissues

Prostate specimens were dissected and cultured as patient-derived explants for 48 hours in the presence and absence of enzalutamide (ENZ; Selleckchem; 10 or 50μM) or the ACC1/2 inhibitor PF-05175157 (Pfizer; 50μM), as previously described (49). In each case, following culture, tissue was either formalin-fixed and paraffin embedded for histology or snap frozen for lipidomics and/or RNA extraction.

### MALDI mass spectrometry imaging

Frozen sections of prostate tissue (10μm) were thaw-mounted on super-frost ultraplus microscope slides and matrix (10mg/ml α-CHCA in methanol) applied to tissue sections by sublimation. The sections were analyzed on a MALDI SYNAPT HDMS Mass Spectrometer (Waters Corporation, Manchester, UK). The laser raster-size was set at 60μm (x,y) and off-tissue areas provided QC spots for data filtering. MALDI raw spectrum files were converted to MSI data files by high definition imaging (HDI) software (Waters Corporation, Manchester, UK). The data processing settings were resolution 8,000 full-width half-height (FWHM) at mass window of 0.02 Da at a (restricted) mass range of *m/z* 400-990 Da. The top 1000 mass features were selected for processing and statistical analysis was carried out using the web-based *MetaboAnalyst R* package (48). The MSI ion map were overlaid or aligned to adjacent pathologically annotated histopathology images to identify morphological regions of interest (ROIs). Equivalent number of data points (3 pixels/mass spectra) from multifocal adenocarcinoma areas and off-tissue regions were selected. The data points were then exported as a single .csv file in the form of ROI’s vs mass features with relative intensity (abundance) as the variable. Before import into *MetaboAnalyst*, the data was filtered in R studio 3.4.4 using baseline packages. The pre-filtered .csv files were uploaded into *MetaboAnalyst R* in the format of spectral bins, data filtering was done by inter-quantile range and data was normalized by log transformation and pareto scaling. Heatmaps were generated by hierarchical clustering and top 25 mass features (identified by ANOVA) were visualized. Principal component analysis (PCA) score plots indicated the relative variation of multifocal ROI’s based on distribution of lipid species. For negative ion mode imaging, the methodology was adapted as follows: Norharmane matrix (7 mg/mL) (Sigma-Aldrich) in CHCl3:MeOH (7:3 v/v) was applied to the tissue sections with a SunCollect sprayer (SunChrom, Friedrichsdorf, Germany). Data processing settings were resolution 8,000 FWHM at mass window of 0.02 Da at a (restricted) mass range of *m/z* 50-990 Da. Full-scan MSI data was imported into Python and a custom script used for ROI selection. All images, mass spectra and boxplots were generated after total ion current normalization of the acquired data.

For validation of ESI-MS/MS data in patient-derived explants, we conducted MALDI imaging using a timsTOF FleX mass spectrometer (Bruker Daltonik, Bremen, Germany). Briefly 10 μm thick frozen tissue sections were thaw mounted onto ITO slides, spray coated with *ca* 500 μL of 7 mg/mL αCHCA matrix using a SunCollect MALDI sprayer (Sunchrom GmbH, Friedrichsdorf, Germany). Data were acquired with a 20 μm pixel size and a 20 μm laser step-size over *m/z* 50-1250. Imaging data was imported into SCiLS Lab 2020a (Bruker Daltonik, Bremen, Germany). Spectra were normalised to total ion count, with weak denoising, and segmentation analysis was preformed using bisecting k-means algorithm with correlation distance metric in SCiLS Lab. Segments which aligned with the epithelia in matched H&E stained sections were selected for each tissue. Box plots of the relative intensity of *m/z* of interest were generated from these segments.

### ESI-MS/MS-based lipidomics

Lipid extracts were generated by homogenizing approximately 40mg of tissue in 800μl PBS with a Dounce or a Precellys (Bertin Technologies) homogenizer. An aliquot of 100μl was set aside for DNA quantification. The remaining 700μl was transferred to a glass tube with Teflon liner and 900μl 1N HCl:CH_3_OH 1:8 (v/v), 800μl CHCl_3_ and 500μg of the antioxidant 2,6-di-tert-butyl-4-methylphenol (BHT) (Sigma, St. Louis, MO) were added. DNA concentration was measured using Hoechst 33258 reagent (Calbiochem, La Jolla, CA). The appropriate lipid standards (Avanti Polar Lipids Inc., Alabaster, AL) were added based on the amount of DNA of the original sample (per mg DNA: 150nmol PC26:0; 50nmol PC28:0; 150nmol PC40:0; 75nmol PE28:0; 8.61nmol PI25:0 and 3nmol PS28:0). After mixing for 5 min in a rotary shaker and phase separation (by centrifugation at 17300xg, for 5 min at 4°C), the lower organic fraction was collected using a glass Pasteur pipette and evaporated using a Savant Speedvac spd111v (Thermo Fisher Scientific, Waltham, MA). The remaining lipid pellet was covered with argon gas and stored at −20 °C. Before ESI-MS/MS measurement, lipid pellets were reconstituted in diluent (CH_3_OH:CHCl_3_:NH_4_OH; 90:10:1.25, v/v/v) according to the amount of DNA in the original cell sample (1μl diluent / 1μg DNA). Phospholipid species were analyzed by ESI-MS/MS on a hybrid quadrupole linear ion trap mass spectrometer (4000 QTRAP system; Applied Biosystems, Foster City, CA) equipped with an Advion TriVersa robotic nanosource for automated sample injection (Advion Biosciences). For quantification of individual phospholipid species, the system was operated in multiple reaction monitoring (MRM) mode. MRM transitions were built based on the release of the phospholipid head group as ion or as neutral species during tandem MS experiments. Analysis was performed using Rapid Lipid Profiling v2.2. Data were corrected for carbon isotope effects. Blank samples consisting of only diluent were measured to determine background signals. Only phospholipid species with an intensity > 5-fold the intensity of the blank were considered true signals. PL were annotated using “lipid subclass” and the “C followed by the total fatty acyl chain length:total number of unsaturated bonds”. The circle plot of lipidomic alterations was generated using the circlize R package (50).

### RNA extraction and sequencing

RNA was extracted from cultured PDE tissues as previously described (51). Total RNA samples were treated with Ribo-Zero to deplete rRNA prior to library construction with the Illumina TruSeq RNA kit. Sequencing was performed on an Illumina NextSeq 500 to generate 1×100bp single-end reads. Library preparation and sequencing were performed at the Genomics Facility of the South Australian Health and Medical Research Institute (Adelaide, Australia). The quality and number of reads for each sample were assessed with FastQC v0.11.3. Adaptors were trimmed from reads, and low-quality bases, with Phred scores < 28, were trimmed from ends of reads, using Trimgalore v0.4.4. Trimmed reads of <20 nucleotides were discarded. Reads passing all quality control steps were aligned to the hg38 assembly of the human genome using TopHat v2.1.1 (52) allowing for up to two mismatches. Reads not uniquely aligned to the genome were discarded. HTSeq-count v0.6.1 (53) was used with the union model to assign uniquely aligned reads to Ensembl Hg38.86-annotated genes. Data were normalized across libraries by the trimmed mean of M-values (TMM) normalization method, implemented in the R v3.5.0, using Bioconductor v3.6 EdgeR v3.20.9 package (54). Only genes expressed at a count-per-million above 0.5 for at least 18 out of 36 samples were analysed for evidence of differential gene expression. Differentially expressed genes were identified using the quasi-likelihood negative binomial generalized log-linear model implemented in EdgeR and were defined as having an FDR adjusted *P* value of < 0.05. Gene Set Enrichment Analysis (GSEA) was undertaken using camera() function in the R limma v3.34.9 package.

### Immunohistochemical Staining

Sections (3μm) of paraffin-embedded cultured patient-derived explants were immunostained essentially as described previously (49). Briefly, antigen retrieval was performed using Tris-EDTA buffer, pH 6.5 using a Biocare Medical Nexgen decloaker at 115°C for 15 min. Tissue slides were then incubated at room temperature with 10% goat serum block. Primary antibody against Ki67 (DAKO, M7240; 1:200), ERG (Abcam, ab92513; 1:400) or pACC1 (Cell Signaling, 3661S; 1:400) was applied and slides incubated overnight at 4°C. Secondary antibody anti-rabbit (DAKO E0432, Lot #20027287) was applied for one hour, followed by HRP-conjugated streptavidin (DAKO, P0397, Lot #20040879) at 1:500 for one hour and visualization by DAB.

### Statistical Analysis

#### Matched patient tissues

*Cohort A*. In the n=21 sample with matched tumour and disease free tissue a mixed effects regression of the within individual difference in lipid variable adjusting for lipid variable in normal tissue sample and individuals age. A random intercept was included per batch, error variance was allowed to differ with batch, and a compound symmetry correlation structure for errors within batches (R package nlme). Restricted maximum likelihoods of this full model is compared with a reduced model without the error variance and correlation structure and the full model chosen only when 2 times the difference in log likelihoods exceeded the 95th percentile of χ2(df=1)=3.84. The lipid variables consisted of log2 transformed species abundance, saturation group abundance and saturation group mean chain length. Prior to these calculations lipid species abundances in each sample were standardized to the median abundance across all samples (ref Dieterle et al 2006).

#### Unmatched patient tissues

*Cohort B*. In the cohort of n=47 with single tissue samples per individual, for associations with sample malignancy similar mixed effects regression to those described above (same random effects structure) were constructed for each lipid variable as outcome, however the only fixed effect covariate was age. In the subset of 28 individuals with malignant samples associations between lipid variables and serum PSA employed the same mixed effects regression models with sample malignancy replaced by log transformed PSA as a fixed effect and sample Gleason score included as an additional covariate.

#### Proliferation associations

For associations with Ki67 both in Day 0 samples and in vehicle vs ENZ treated samples we analyze repeated count data across fields of view within a sample using a beta-binomial mixed effects regression (R package glmmTMB). In the Day 0 analyses, the lipid variables are the primary predictors of interest with batch and age being included as fixed effect covariates. In the treated samples, the primary predictor of interest is the batch-adjusted difference between ENZ and vehicle, with batch, age and lipid variable vehicle as fixed effects. In all models, a random intercept is included per individual, with a logit link for the mean Ki67 cell positivity prevalence.

For each set of outcomes the distribution of p-values for the species abundance associations is assessed to determine the presence of a signal (deviation from the uniform distribution) (55) and the FDR reported for the number of significant associations defined as p<0.01. To address concerns that an apparent signal may be due to a combination of insufficient comparisons and clustering between lipid species, we perform a permutation test with 1000 permutations of the clinical outcome, and define the fraction of significant associations are beyond that observed in the original cohort as the permutation p-value. These permutation analyses are performed within batches and where appropriate within malignancy groups, and by design retain the between lipid correlation structure.

#### Other statistical analyses

Statistical analysis for lipid or immunostaining quantification and PDE *ex vivo* culture experiments, was carried out using GraphPad Prism software v7.02 (2016, GraphPad Software). Significance was measured by two-tailed unpaired t-test or oneway ANOVA with Dunnett’s multiple comparison test as indicated. Significance is expressed as \**P* < 0.05, \*\**P* < 0.01 and \*\*\**P* < 0.001.

## Acknowledgments and Grant Support

This research was supported by The Movember Foundation/Prostate Cancer Foundation of Australia (MRTA3); The Prostate Cancer Foundation of Australia (ID NDDA); The National Health and Medical Research Council (NHMRC) of Australia (ID APP1121057, APP1145777, APP1138242, APP1130077); a Movember/NBCF Collaborative Linkage Grant (MNBCF-17-012), KU Leuven grants C1 (C16/15/073) and C3 (C32/17/052); Research Foundation-Flanders (FWO) grants G.841.15 and G0E0817N; Interreq V-A EMR23 EURLIPIDS; Stichting tegen Kanker; Kom op tegen Kanker; and the Fondation Fournier-Majoie pour l’Innovation. L.M.B. was supported by a Future Fellowship from the Australian Research Council (FT130101004), and she and L.A.S are supported by Principal Cancer Research Fellowships produced with the financial and other support of Cancer Council SA’s Beat Cancer Project on behalf of its donors and the State Government of South Australia through the Department of Health. C.Y.M. is supported by a PhD Scholarship from the University of Adelaide and the Freemason’s Foundation Centre for Men’s Health at the University of Adelaide; Z.D.N. is supported by an Early Career Fellowship from the National Health and Medical Research Council of Australia (ID 1138648); X.S. is supported by a Fellowship from the Research Foundation Flanders (FWO); D.J.L. is supported by an EMBL Australia group leader award.

## Author Contributions

Conception and design: LMB, MMC, JVS.

Development of methodology: LMB, MMC, JM, SM, XS, JD, RD, EW, PJT, MFS, JVS.

Pathology of patient samples: RD, JS, JK, DG.

Acquisition of data: LMB, MMC, CYM, JM, SI, SM, XS, FV, KB, RD.

Analysis and interpretation of data: LMB, CYM, ADV, DW, MM, JD, ZDN, LAS, PJT, MFS, DJL, WDT, LGH, MMC, JVS.

Study Supervision: LMB, PJT, MFS, DJL, MMC, JVS.

Drafting of the manuscript: LMB, MMC, JVS.

Revision of the manuscript: All authors read and reviewed the manuscript

## Supplementary Figure Legends

**Supplementary Figure 1.**
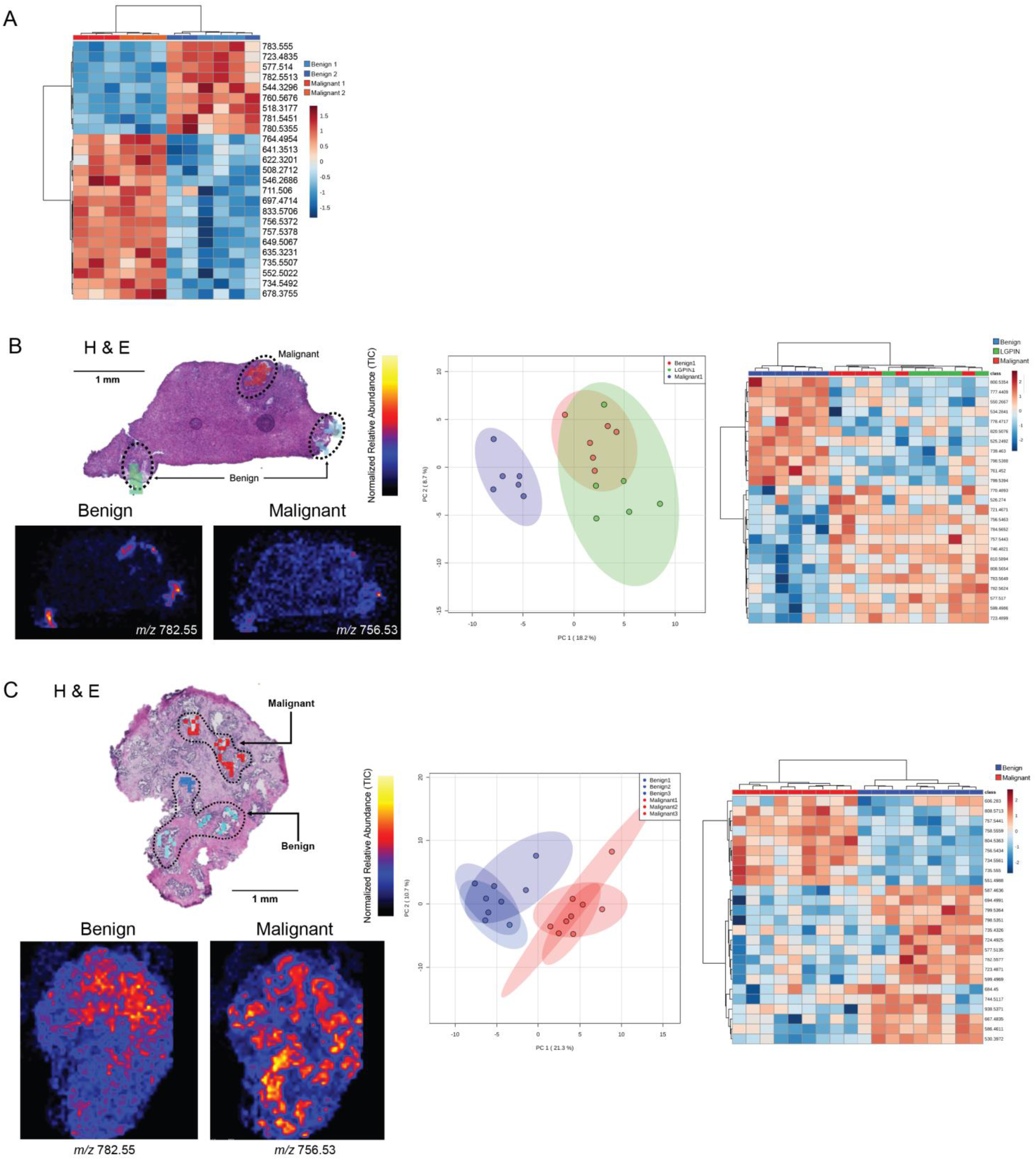
MALDI-mass spectrometry imaging of regions of interest in pathologically heterogeneous prostate tissue from 3 individual prostate cancer patients, including ion maps of two representative examples of histology-restricted lipid masses, principal component and heatmap analysis of the top 25 mass features distinguishing benign from malignant regions of tissue for each patient.

**Supplementary Figure 2.**
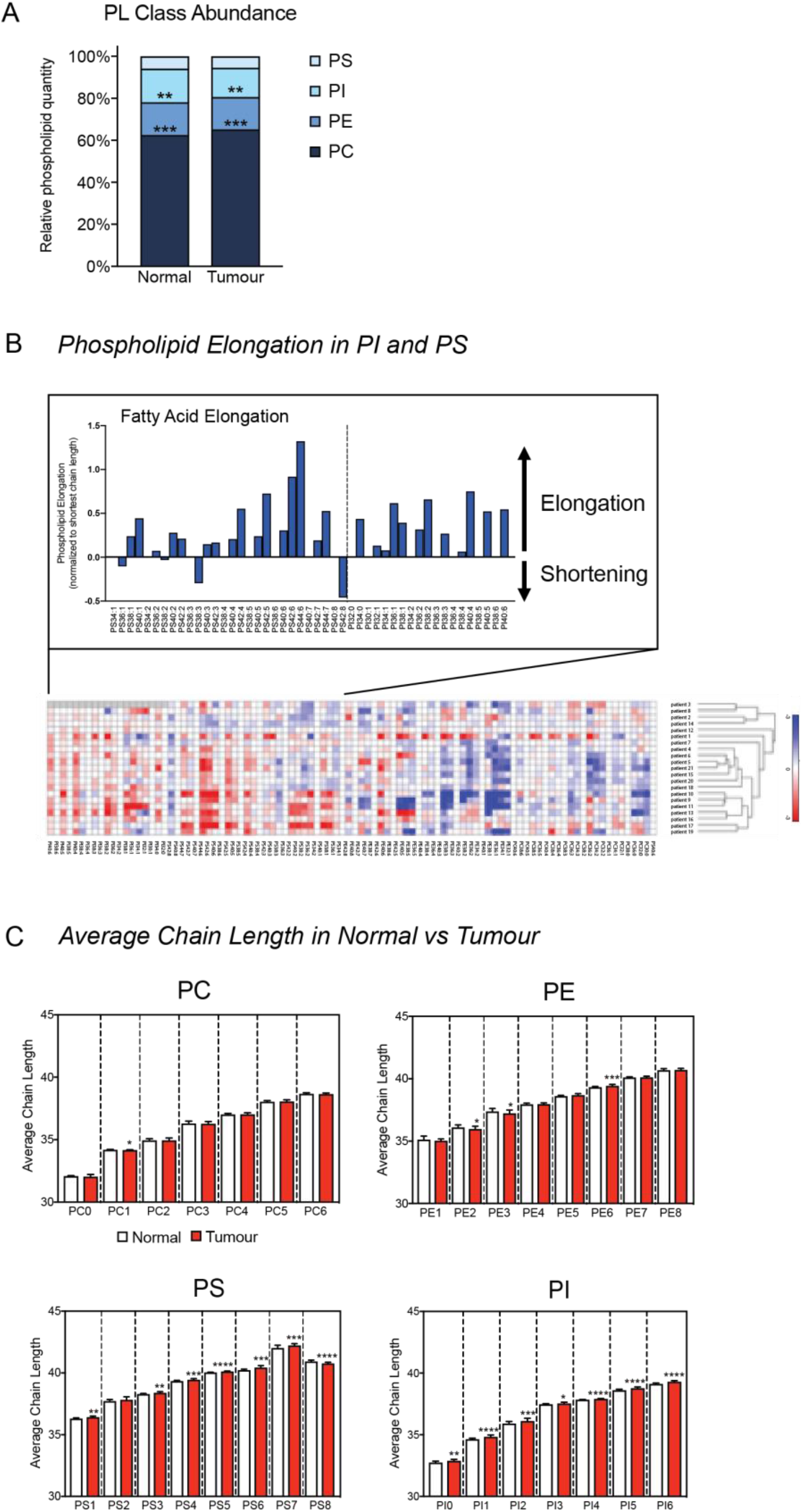
Cancer-related changes in phospholipid composition in matched non-malignant and malignant tissues from prostate cancer patients (n=21). **A.** Relative proportions of PC, PE, PS and PI phospholipid classes across the patient cohort. **B.** Heatmap clustering of changes in PI, PS, PE and PC fatty acid chain length and saturation in prostate tumours versus matched non-malignant tissue. Each row represents a patient, and each column represents a different phospholipid species. The elongation can be observed as vertical striation patterns of increased chain lengths for each saturation group, and is represented graphically for the PS and PI species in the inset box. **C.** Tumor-related changes in average chain length for each main phospholipid class.

**Supplementary Figure 3.**
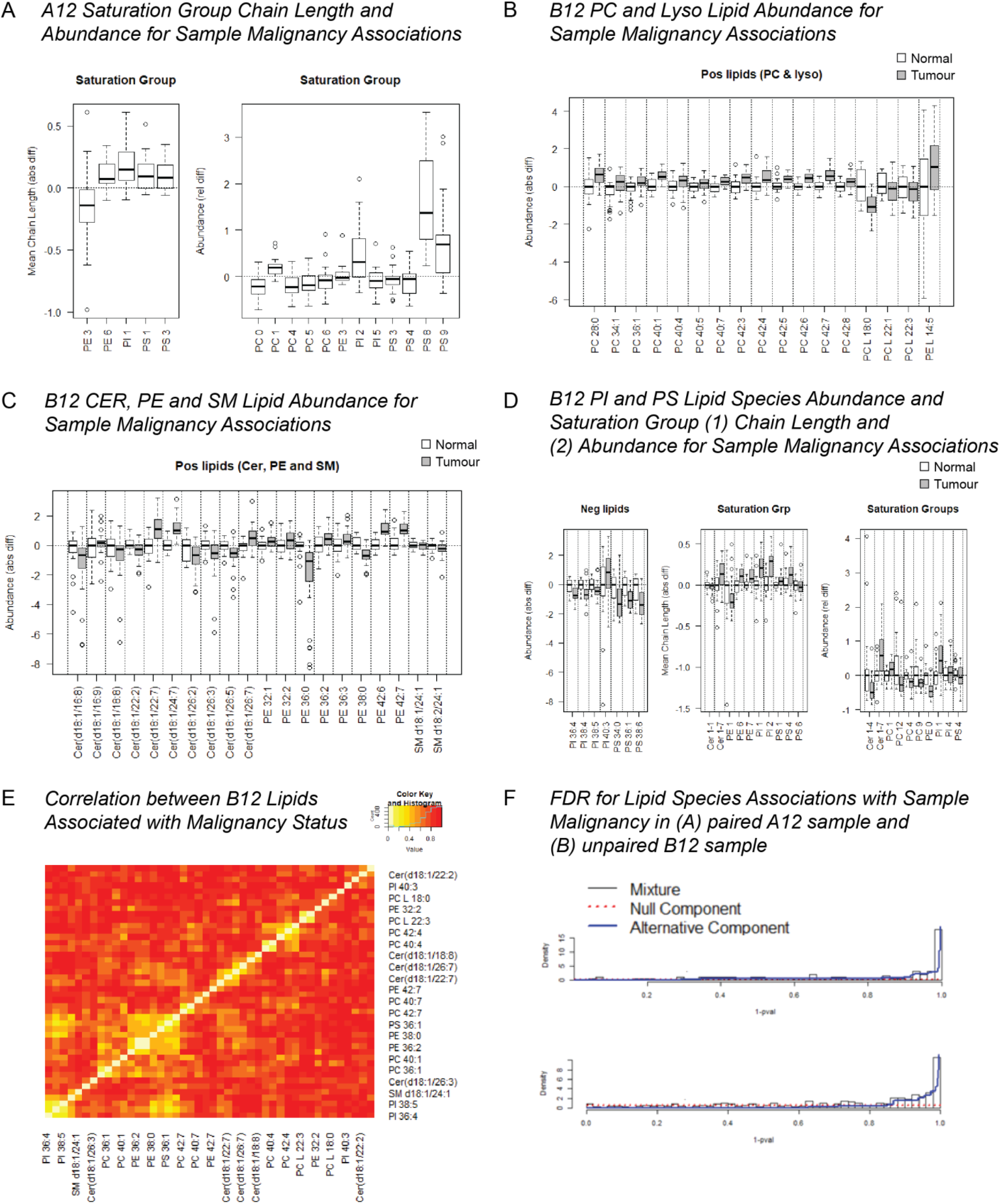
Phospholipid species associations with malignancy. **A.** FDR for PL associations with malignancy in Cohorts A and B. **B.** Saturation group chain length and abundance associations with malignancy in Cohort A. **C, D, E.** PL abundance associations with malignancy in Cohort B. **F.** Correlation plot of individual phospholipid species significantly associated with sample malignancy in Cohort B.

**Supplementary Figure 4.**
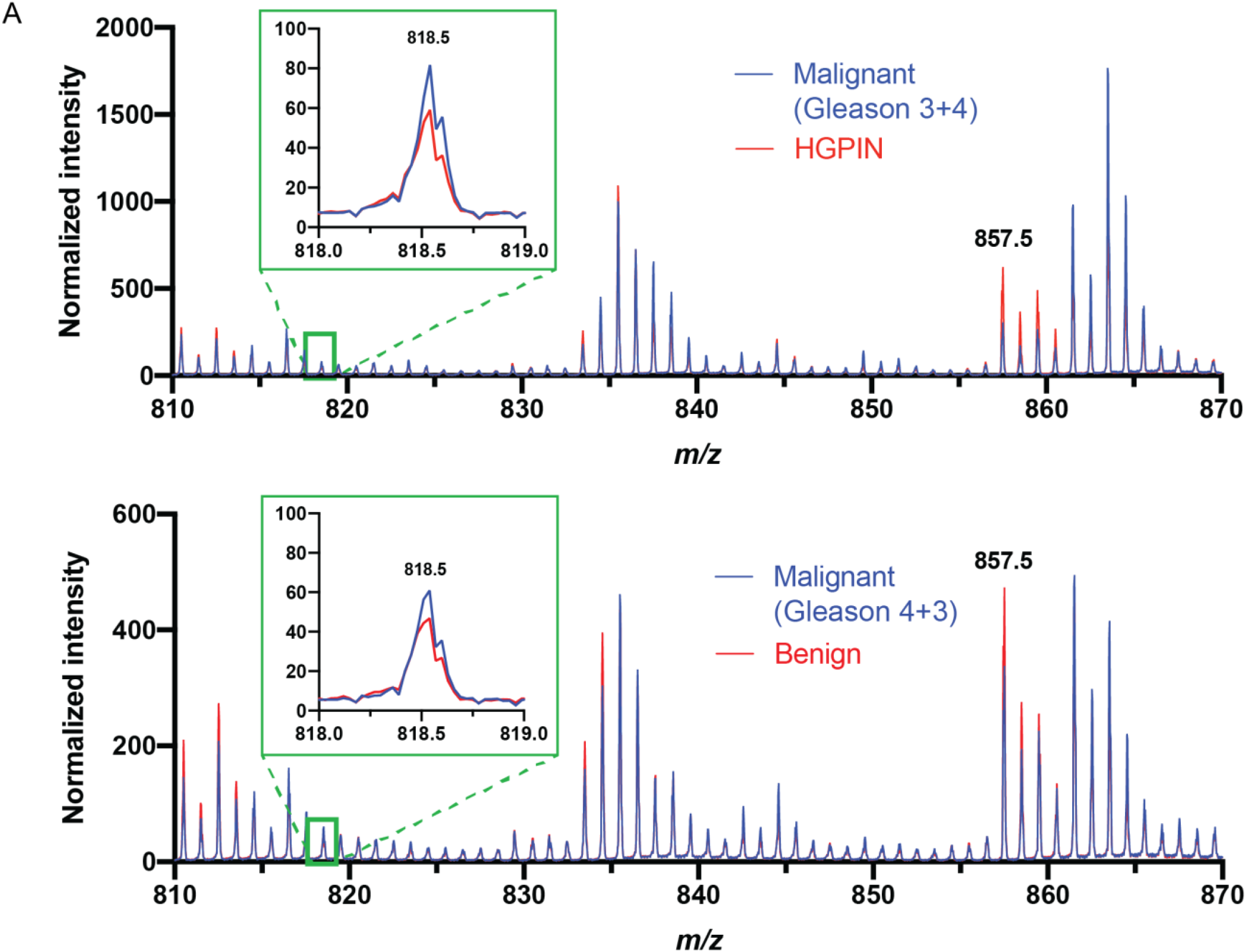
MALDI-mass spectrometry imaging-derived spectra of discrete histological foci (non-malignant versus malignant as indicated) in 2 independent prostate cancer patients.

**Supplementary Figure 5.**
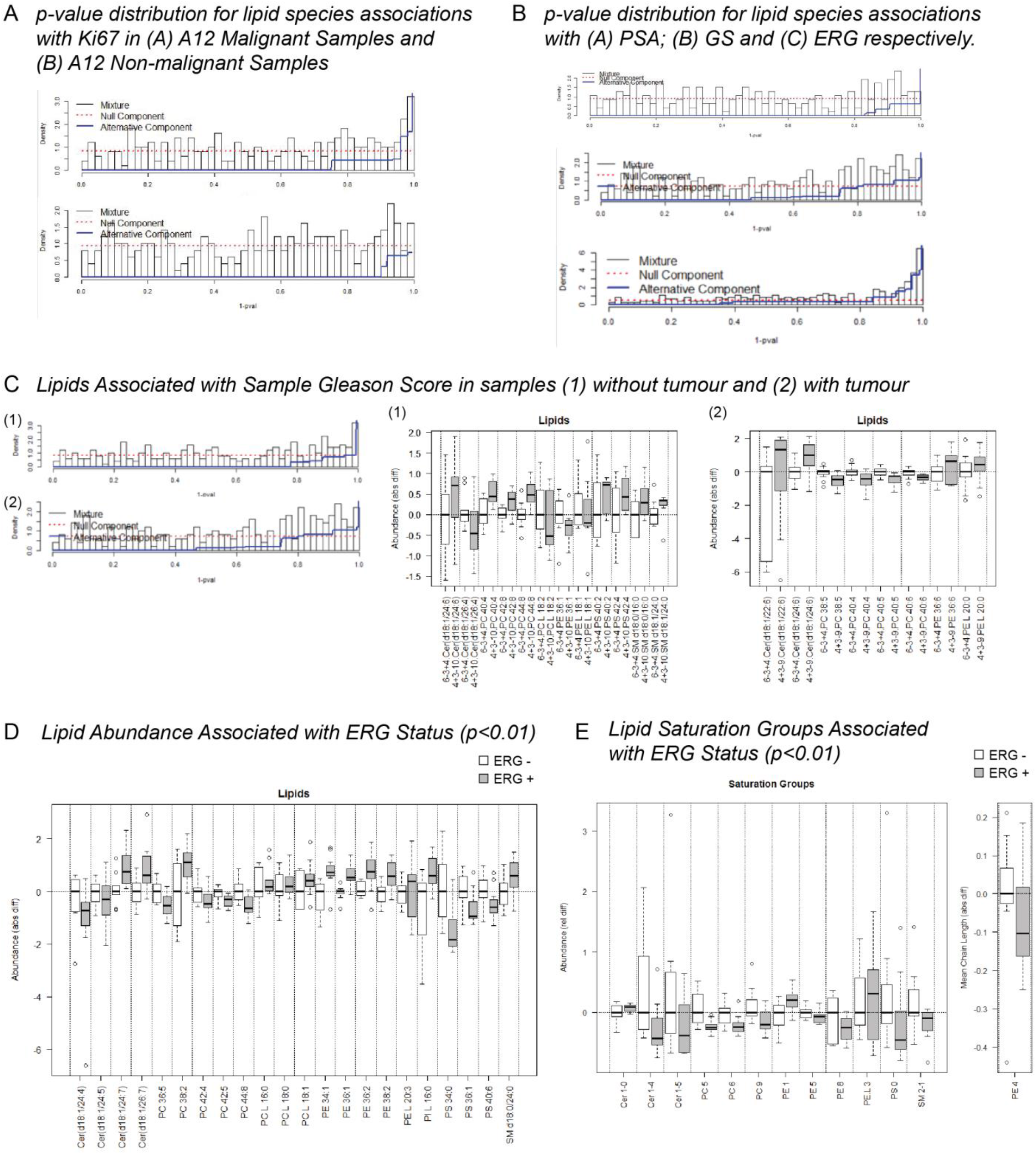
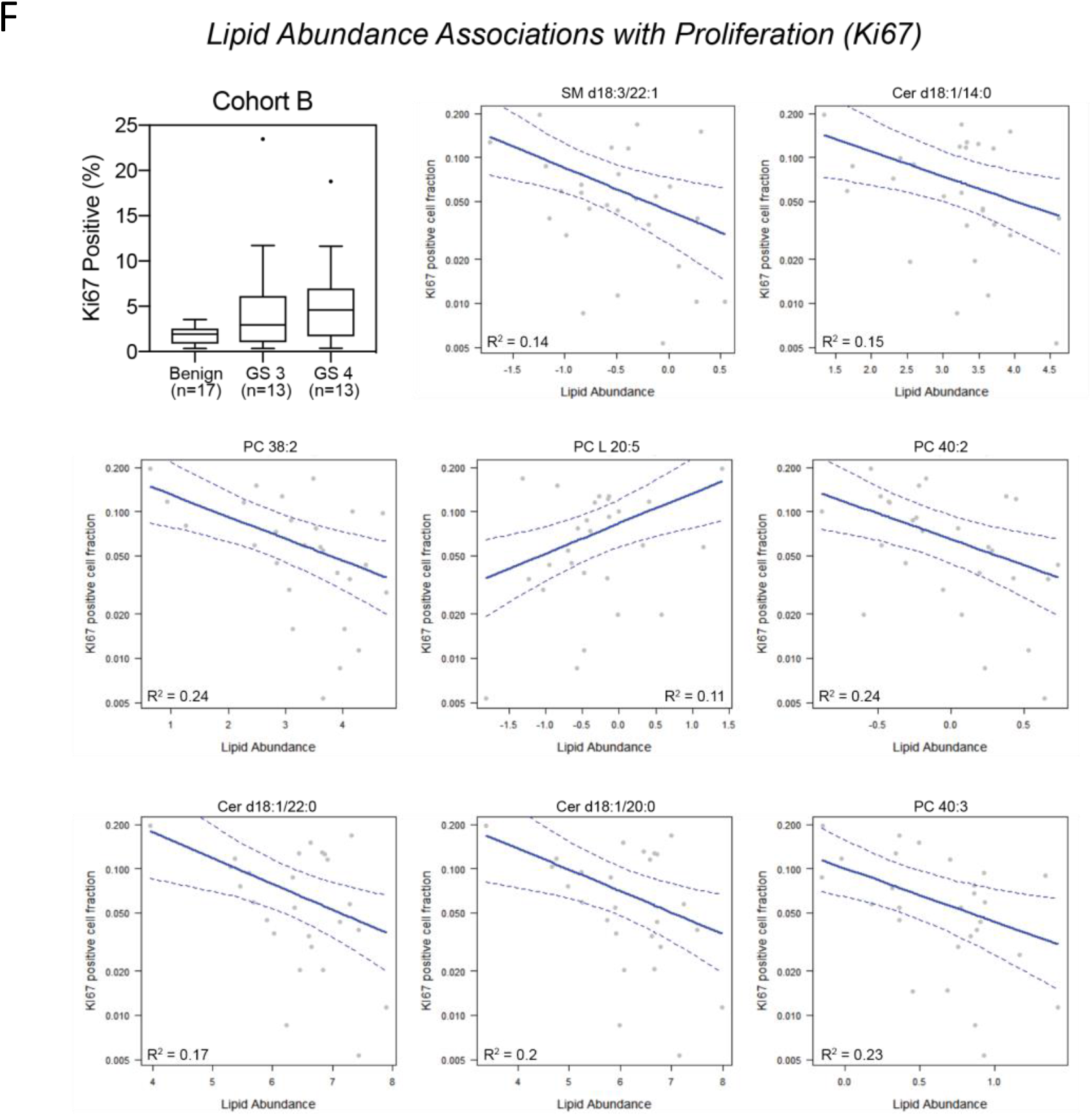
Associations of lipid measures with tumor clinicopathological features. p-value distribution for associations of lipid species with **A.** Ki67 proliferative index, and **B.** serum prostate specific antigen levels, tumor Gleason score, and ERG positivity. **C.** PL abundance associations with original tumor pathology. **D, E.** PL associations with ERG positivity status. **F.** PL associations with Ki67 positivity.

**Supplementary Figure 6.**
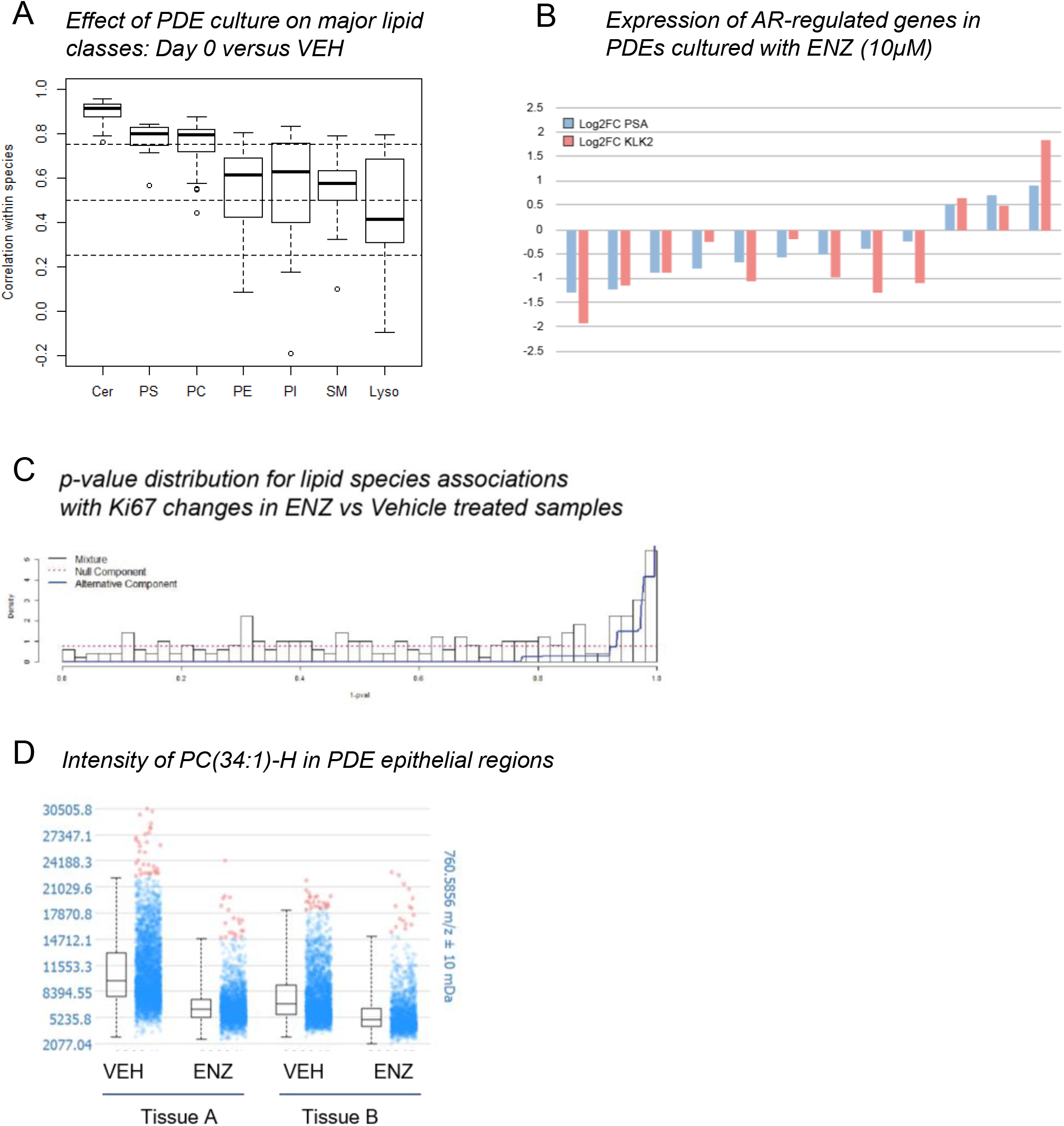
Associations of lipid measures with response to the AR antagonist enzalutamide in patient-derived explants of clinical prostate tissues. **A.** Effect of ex vivo culture on PL profiles in prostate tissues, demonstrated by correlations in lipids between uncultured and vehicle-cultured samples. **B.** Treatment-related changes in mRNA expression of the AR-target genes, *KLK3* (prostate specific antigen) and *KLK2*. **C.** P-value distribution for PL associations with proliferative response to enzalutamide. **D.** Box plots of normalized ion intensity for PC34:1-H+ in epithelial regions of enzalutamide-treated versus vehicle-treated explants from two separate tissue cores.

**Supplementary Figure 7.**
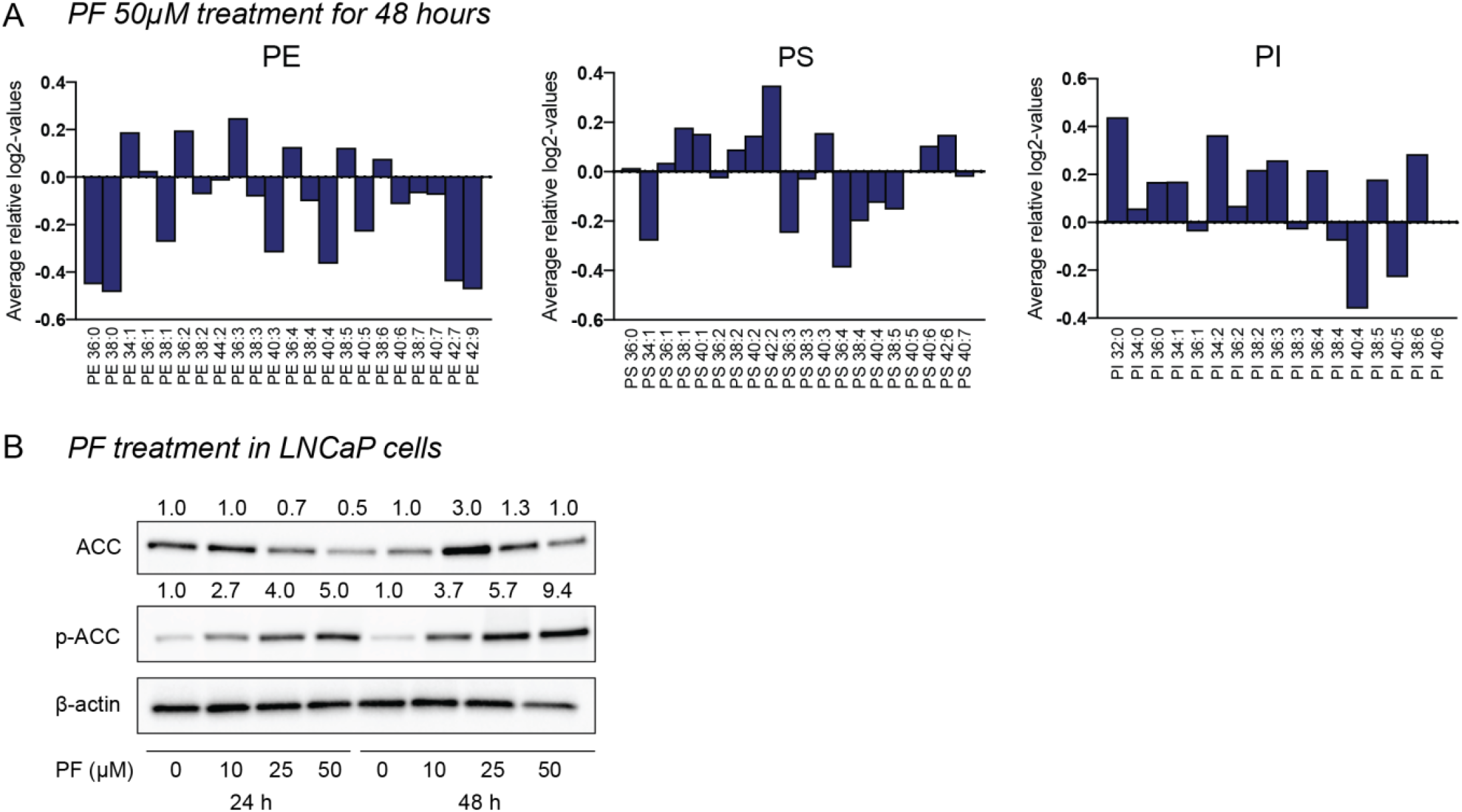
**A.** Altered fatty acyl chain length for PE, PS and PI lipids in patient-derived explants cultured in the ACC inhibitor PF-05175157 compared to vehicle control. **B.** Dose-dependent increase in pACC1/ACC1 levels in LNCaP prostate cancer cells cultured with PF-05175157 (50μM) for 24 or 48 hours.

## References

1. Bray F, Ferlay J, Soerjomataram I, Siegel RL, Torre LA, Jemal A. Global cancer statistics 2018: GLOBOCAN estimates of incidence and mortality worldwide for 36 cancers in 185 countries. CA Cancer J Clin 2018;68(6):394–424.

2. Butler LM, Perone Y, Dehairs J, Lupien LE, de Laat V, Talebi A, et al. Lipids and cancer: Emerging roles in pathogenesis, diagnosis and therapeutic intervention. Adv Drug Deliv Rev 2020.

3. Hanahan D, Weinberg RA. Hallmarks of cancer: the next generation. Cell 2011;144(5):646–74.

4. Koundouros N, Poulogiannis G. Reprogramming of fatty acid metabolism in cancer. Br J Cancer 2020;122(1):4–22.

5. Bader DA, McGuire SE. Tumour metabolism and its unique properties in prostate adenocarcinoma. Nat Rev Urol 2020;17(4):214–31.

6. Zadra G, Loda M. Metabolic Vulnerabilities of Prostate Cancer: Diagnostic and Therapeutic Opportunities. Cold Spring Harb Perspect Med 2018;8(10).

7. Butler LM, Centenera MM, Swinnen JV. Androgen control of lipid metabolism in prostate cancer: novel insights and future applications. Endocr Relat Cancer 2016;23(5):R219–27.

8. Crowe FL, Appleby PN, Travis RC, Barnett M, Brasky TM, Bueno-de-Mesquita HB, et al. Circulating fatty acids and prostate cancer risk: individual participant metaanalysis of prospective studies. J Natl Cancer Inst 2014;106(9).

9. Patel N, Vogel R, Chandra-Kuntal K, Glasgow W, Kelavkar U. A novel three serum phospholipid panel differentiates normal individuals from those with prostate cancer. PLoS One 2014;9(3):e88841.

10. Lin HM, Mahon KL, Weir JM, Mundra PA, Spielman C, Briscoe K, et al. A distinct plasma lipid signature associated with poor prognosis in castration-resistant prostate cancer. Int J Cancer 2017;141(10):2112–20.

11. Rysman E, Brusselmans K, Scheys K, Timmermans L, Derua R, Munck S, et al. De novo lipogenesis protects cancer cells from free radicals and chemotherapeutics by promoting membrane lipid saturation. Cancer Res 2010;70(20):8117–26.

12. Sorvina A, Bader CA, Caporale C, Carter EA, Johnson IRD, Parkinson-Lawrence EJ, et al. Lipid profiles of prostate cancer cells. Oncotarget 2018;9(85):35541–52.

13. Burch TC, Isaac G, Booher CL, Rhim JS, Rainville P, Langridge J, et al. Comparative Metabolomic and Lipidomic Analysis of Phenotype Stratified Prostate Cells. PLoS One 2015;10(8):e0134206.

14. Goto T, Terada N, Inoue T, Kobayashi T, Nakayama K, Okada Y, et al. Decreased expression of lysophosphatidylcholine (16:0/OH) in high resolution imaging mass spectrometry independently predicts biochemical recurrence after surgical treatment for prostate cancer. Prostate 2015;75(16):1821–30.

15. Goto T, Terada N, Inoue T, Nakayama K, Okada Y, Yoshikawa T, et al. The expression profile of phosphatidylinositol in high spatial resolution imaging mass spectrometry as a potential biomarker for prostate cancer. PLoS One 2014;9(2):e90242.

16. Morse N, Jamaspishvili T, Simon D, Patel PG, Ren KYM, Wang J, et al. Reliable identification of prostate cancer using mass spectrometry metabolomic imaging in needle core biopsies. Laboratory Investigation 2019.

17. Randall EC, Zadra G, Chetta P, Lopez BGC, Syamala S, Basu SS, et al. Molecular Characterization of Prostate Cancer with Associated Gleason Score Using Mass Spectrometry Imaging. Mol Cancer Res 2019;17(5):1155–65.

18. Beer TM, Armstrong AJ, Rathkopf DE, Loriot Y, Sternberg CN, Higano CS, et al. Enzalutamide in metastatic prostate cancer before chemotherapy. N Engl J Med 2014;371(5):424–33.

19. Scher HI, Fizazi K, Saad F, Taplin ME, Sternberg CN, Miller K, et al. Increased survival with enzalutamide in prostate cancer after chemotherapy. N Engl J Med 2012;367(13):1187–97.

20. Griffith DA, Kung DW, Esler WP, Amor PA, Bagley SW, Beysen C, et al. Decreasing the rate of metabolic ketone reduction in the discovery of a clinical acetyl-CoA carboxylase inhibitor for the treatment of diabetes. J Med Chem 2014;57(24):10512–26.

21. Zhang AY, Chiam K, Haupt Y, Fox S, Birch S, Tilley W, et al. An analysis of a multiple biomarker panel to better predict prostate cancer metastasis after radical prostatectomy. Int J Cancer 2019;144(5):1151–59.

22. Bandu R, Mok HJ, Kim KP. Phospholipids as cancer biomarkers: Mass spectrometrybased analysis. Mass Spectrom Rev 2018;37(2):107–38.

23. Naguib A, Bencze G, Engle DD, Chio, II, Herzka T, Watrud K, et al. p53 mutations change phosphatidylinositol acyl chain composition. Cell Rep 2015;10(1):8–19.

24. Falvella FS, Pascale RM, Gariboldi M, Manenti G, De Miglio MR, Simile MM, et al. Stearoyl-CoA desaturase 1 (Scd1) gene overexpression is associated with genetic predisposition to hepatocarcinogenesis in mice and rats. Carcinogenesis 2002;23(11):1933–6.

25. Fritz V, Benfodda Z, Rodier G, Henriquet C, Iborra F, Avances C, et al. Abrogation of de novo lipogenesis by stearoyl-CoA desaturase 1 inhibition interferes with oncogenic signaling and blocks prostate cancer progression in mice. Mol Cancer Ther 2010;9(6):1740–54.

26. Griffitts J, Tesiram Y, Reid GE, Saunders D, Floyd RA, Towner RA. In vivo MRS assessment of altered fatty acyl unsaturation in liver tumor formation of a TGF alpha/c-myc transgenic mouse model. J Lipid Res 2009;50(4):611–22.

27. Li J, Ding SF, Habib NA, Fermor BF, Wood CB, Gilmour RS. Partial characterization of a cDNA for human stearoyl-CoA desaturase and changes in its mRNA expression in some normal and malignant tissues. Int J Cancer 1994;57(3):348–52.

28. Romanuik TL, Ueda T, Le N, Haile S, Yong TM, Thomson T, et al. Novel biomarkers for prostate cancer including noncoding transcripts. Am J Pathol 2009;175(6):2264–76.

29. Tamura K, Makino A, Hullin-Matsuda F, Kobayashi T, Furihata M, Chung S, et al. Novel lipogenic enzyme ELOVL7 is involved in prostate cancer growth through saturated long-chain fatty acid metabolism. Cancer Res 2009;69(20):8133–40.

30. Wang HW, Wu YH, Hsieh JY, Liang ML, Chao ME, Liu DJ, et al. Pediatric primary central nervous system germ cell tumors of different prognosis groups show characteristic miRNome traits and chromosome copy number variations. BMC Genomics 2010;11:132.

31. Marien E, Meister M, Muley T, Fieuws S, Bordel S, Derua R, et al. Non-small cell lung cancer is characterized by dramatic changes in phospholipid profiles. Int J Cancer 2015;137(7):1539–48.

32. Swinnen JV, Vanderhoydonc F, Elgamal AA, Eelen M, Vercaeren I, Joniau S, et al. Selective activation of the fatty acid synthesis pathway in human prostate cancer. Int J Cancer 2000;88(2):176–9.

33. Swinnen JV, Heemers H, Deboel L, Foufelle F, Heyns W, Verhoeven G. Stimulation of tumor-associated fatty acid synthase expression by growth factor activation of the sterol regulatory element-binding protein pathway. Oncogene 2000;19(45):5173–81.

34. Van de Sande T, De Schrijver E, Heyns W, Verhoeven G, Swinnen JV. Role of the phosphatidylinositol 3’-kinase/PTEN/Akt kinase pathway in the overexpression of fatty acid synthase in LNCaP prostate cancer cells. Cancer Res 2002;62(3):642–6.

35. Moreau K, Dizin E, Ray H, Luquain C, Lefai E, Foufelle F, et al. BRCA1 affects lipid synthesis through its interaction with acetyl-CoA carboxylase. J Biol Chem 2006;281(6):3172–81.

36. Kinlaw WB, Church JL, Harmon J, Mariash CN. Direct evidence for a role of the “spot 14” protein in the regulation of lipid synthesis. J Biol Chem 1995;270(28):16615–8.

37. Sabbisetti V, Di Napoli A, Seeley A, Amato AM, O’Regan E, Ghebremichael M, et al. p63 promotes cell survival through fatty acid synthase. PLoS One 2009;4(6):e5877.

38. Swinnen JV, Ulrix W, Heyns W, Verhoeven G. Coordinate regulation of lipogenic gene expression by androgens: evidence for a cascade mechanism involving sterol regulatory element binding proteins. Proceedings of the National Academy of Sciences of the United States of America 1997;94(24):12975–80.

39. Harayama T, Riezman H. Understanding the diversity of membrane lipid composition. Nat Rev Mol Cell Biol 2018;19(5):281–96.

40. Rogers KR, Kikawa KD, Mouradian M, Hernandez K, McKinnon KM, Ahwah SM, et al. Docosahexaenoic acid alters epidermal growth factor receptor-related signaling by disrupting its lipid raft association. Carcinogenesis 2010;31(9):1523–30.

41. Gillet L, Roger S, Bougnoux P, Le Guennec JY, Besson P. Beneficial effects of omega-3 long-chain fatty acids in breast cancer and cardiovascular diseases: voltage-gated sodium channels as a common feature? Biochimie 2011;93(1):4–6.

42. Jude S, Roger S, Martel E, Besson P, Richard S, Bougnoux P, et al. Dietary long-chain omega-3 fatty acids of marine origin: a comparison of their protective effects on coronary heart disease and breast cancers. Prog Biophys Mol Biol 2006;90(1-3):299–325.

43. Kiebish MA, Han X, Cheng H, Chuang JH, Seyfried TN. Cardiolipin and electron transport chain abnormalities in mouse brain tumor mitochondria: lipidomic evidence supporting the Warburg theory of cancer. Journal of lipid research 2008;49(12):2545–56.

44. Hilvo M, Denkert C, Lehtinen L, Muller B, Brockmoller S, Seppanen-Laakso T, et al. Novel theranostic opportunities offered by characterization of altered membrane lipid metabolism in breast cancer progression. Cancer Res 2011;71(9):3236–45.

45. Beckers A, Organe S, Timmermans L, Scheys K, Peeters A, Brusselmans K, et al. Chemical inhibition of acetyl-CoA carboxylase induces growth arrest and cytotoxicity selectively in cancer cells. Cancer Res 2007;67(17):8180–7.

46. Brusselmans K, De Schrijver E, Verhoeven G, Swinnen JV. RNA interference-mediated silencing of the acetyl-CoA-carboxylase-alpha gene induces growth inhibition and apoptosis of prostate cancer cells. Cancer Res 2005;65(15):6719–25.

47. Peck B, Schug ZT, Zhang Q, Dankworth B, Jones DT, Smethurst E, et al. Inhibition of fatty acid desaturation is detrimental to cancer cell survival in metabolically compromised environments. Cancer Metab 2016;4:6.

48. Waltregny D, Alami Y, Clausse N, de Leval J, Castronovo V. Overexpression of the homeobox gene HOXC8 in human prostate cancer correlates with loss of tumor differentiation. Prostate 2002;50(3):162–9.

49. Centenera MM, Hickey TE, Jindal S, Ryan NK, Ravindranathan P, Mohammed H, et al. A patient-derived explant (PDE) model of hormone-dependent cancer. Mol Oncol 2018;12(9):1608–22.

50. Gu Z, Gu L, Eils R, Schlesner M, Brors B. circlize Implements and enhances circular visualization in R. Bioinformatics 2014;30(19):2811–2.

51. Armstrong HK, Gillis JL, Johnson IRD, Nassar ZD, Moldovan M, Levrier C, et al. Dysregulated fibronectin trafficking by Hsp90 inhibition restricts prostate cancer cell invasion. Sci Rep 2018;8(1):2090.

52. Kim D, Pertea G, Trapnell C, Pimentel H, Kelley R, Salzberg SL. TopHat2: accurate alignment of transcriptomes in the presence of insertions, deletions and gene fusions. Genome Biol 2013;14(4):R36.

53. Anders S, Pyl PT, Huber W. HTSeq--a Python framework to work with high-throughput sequencing data. Bioinformatics 2015;31(2):166–9.

54. Robinson MD, McCarthy DJ, Smyth GK. edgeR: a Bioconductor package for differential expression analysis of digital gene expression data. Bioinformatics 2010;26(1):139–40.

55. Strimmer K. A unified approach to false discovery rate estimation. BMC Bioinformatics 2008;9:303.

